# A trio of brain networks dynamically orchestrates music recognition

**DOI:** 10.64898/2026.11.09.765255

**Authors:** Barbara A. P. Grosjean, Leonardo Bonetti, Mattia Rosso, Meghana Ramineni, Alejandro O. Blenkmann, Tor Endestad, Anne-Kristin Solbakk, Olivia Kim-McManus, Jon T. Willie, Peter Brunner, Mohammad Dastjerdi, Jack J. Lin, Robert T. Knight, David R. Quiroga-Martinez

## Abstract

Recognition memory continuously integrates internal representations with sensory information to guide adaptive behavior. Yet most theoretical models conceptualize recognition as the retrieval of an isolated stimulus through a discrete sequence of steps including sensory processing, hippocampal retrieval, and cortical reinstatement. However, in natural settings, perception, memory, and predictions interact continuously within an evolving context to guide adaptive responses. It is unclear how brain networks coordinate the integration of internal and external information during dynamic recognition. Here, we combined intracranial recordings (N = 30) of high-frequency activity (70–160 Hz) and dimensionality reduction to characterize distributed network dynamics during music recognition. We identified three interacting functional networks supporting sensory processing, sensory-memory integration, and decision formation, whose temporal dynamics predicted the timing of behavior. During recognition of memorized sequences, sensory activity progressively attenuated, sensory inputs were combined with mnemonic and contextual information, and decision-related activity evolved toward sensorimotor engagement. Novel sequences disrupted this organization by sustaining sensory responses and accelerating decision-related activity. Sensory and integrative networks communicated bidirectionally. Our findings reveal recognition as a distributed, context-dependent, and goal-directed process emerging from the continuous and bidirectional coordination of large-scale brain networks.

## 1. Introduction

Recognition memory enables the interpretation of ongoing sensory information based on prior experience to guide adaptive behavior^1,2^. Rather than simply determining whether a stimulus has been encountered before, recognition requires the continuous integration of external sensory evidence with internally generated representations, allowing individuals to interpret events within their context, anticipate future inputs, and update behavior^3,4^. Recognition should therefore be viewed not only as the successful matching of two pieces of information, but as an active inferential process that continuously integrates perception, memory, and expectations to support goal-directed behavior.

Recognition has traditionally been investigated using paradigms based on the retrieval of isolated stimuli. Consequently, dominant theoretical models describe recognition as the association between a sensory cue and its stored representation through mechanisms such as hippocampal pattern completion, recollection, and cortical reinstatement^5–8^. In these models, sensory information propagates through cortical hierarchies toward the medial temporal lobe (MTL), where partial cues trigger the recovery of stored representations, which are subsequently reinstated across distributed cortical regions^6,9,10^. This framework has shaped our understanding of associative memory retrieval, but it conceptualizes recognition as a succession of processing stages, emphasizing the recovery of stored representations rather than their continuous interaction with ongoing perception^11^.

However, increasing evidence suggests that recognition emerges from recurrent exchanges between perceptual and mnemonic systems. The hippocampus may dynamically alternate between receiving sensory inputs and broadcasting mnemonic information depending on behavioral demands^7,9,10^, while mnemonic representations can bias sensory processing before or during perception through pattern completion and contextual predictions^12–15^. These advances have often been established through the study of individual brain regions or isolated circuits, providing a fragmented understanding of how distributed brain systems integrate sensory evidence, memory, and predictions during recognition in naturalistic settings. Consequently, it is necessary to study whole-brain network dynamics during recognition in situations where sensory information unfolds progressively over time, allowing memory, perception, prediction, and adaptive behavior to continuously interact.

Musical sequences provide such a framework as auditory perception inherently unfolds over time^16^ and naturally embeds incoming events within an evolving predictive context. Previous MEG studies characterized large-scale spatiotemporal dynamics during sequence processing and demonstrated the importance of contextual information for perception and memory^17,18^. However, the distributed neural architecture supporting progressive recognition remains incomplete. In particular, the rapid interactions between perceptual, mnemonic, integrative, and decision-making systems cannot be fully resolved with non-invasive recordings because of their limited spatial or temporal resolution.

Here, we combined intracranial stereo-electroencephalography (sEEG) with whole-brain dimensionality reduction to characterize high-frequency activity (HFA) (70–160 Hz) during auditory recognition. HFA provides a particularly informative measure of local cortical processing because it closely reflects population firing and has been consistently associated with memory encoding and retrieval^19–27^. Participants performed a musical recognition task involving previously learned and novel melodic sequences, allowing recognition to unfold progressively within a temporally structured context. We demonstrate that recognition relies on three interacting functional networks: a sensory processing network tracking incoming auditory information, a decision-making network gradually engaging sensorimotor processes, and an integrative network combining sensory and mnemonic information. Rather than operating sequentially, these networks interact continuously through reciprocal bottom-up and top-down exchanges, revealing recognition memory as a distributed, dynamic, and adaptive neurocognitive process.

## 2. Results

### 2.1. Behavior

We investigated how large-scale high-gamma dynamics support ongoing and in-context recognition by analyzing sEEG recordings from 30 patients (16 female; age = 37.32 +/− 11.91 SD; 2,574 electrodes in total) performing an auditory memorized/novel recognition task. Participants first listened to and memorized the first 4 bars of Bach’s Prelude in C minor repeated twice. Then, they completed an auditory recognition task in which they identified five-tone melodic excerpts that were memorized (M) with the prelude or completely novel (N). 24 trials were done per participant for each experimental condition. Participants performed above chance (estimated accuracy = 92.6%; OR = 12.43, 95% CI [10.46, 14.78], z = 28.61, one-sided p <.001) and were more accurate for novel than memorized melodies (OR = 3.88, 95% CI [2.67, 5.65], z = 7.09, p <.001). Reaction times were similar between conditions (β = 0.014, 95% CI [−0.004, 0.033], p =.127) (Fig. 1). Only correct trials were retained for further analysis.

**Figure 1.**
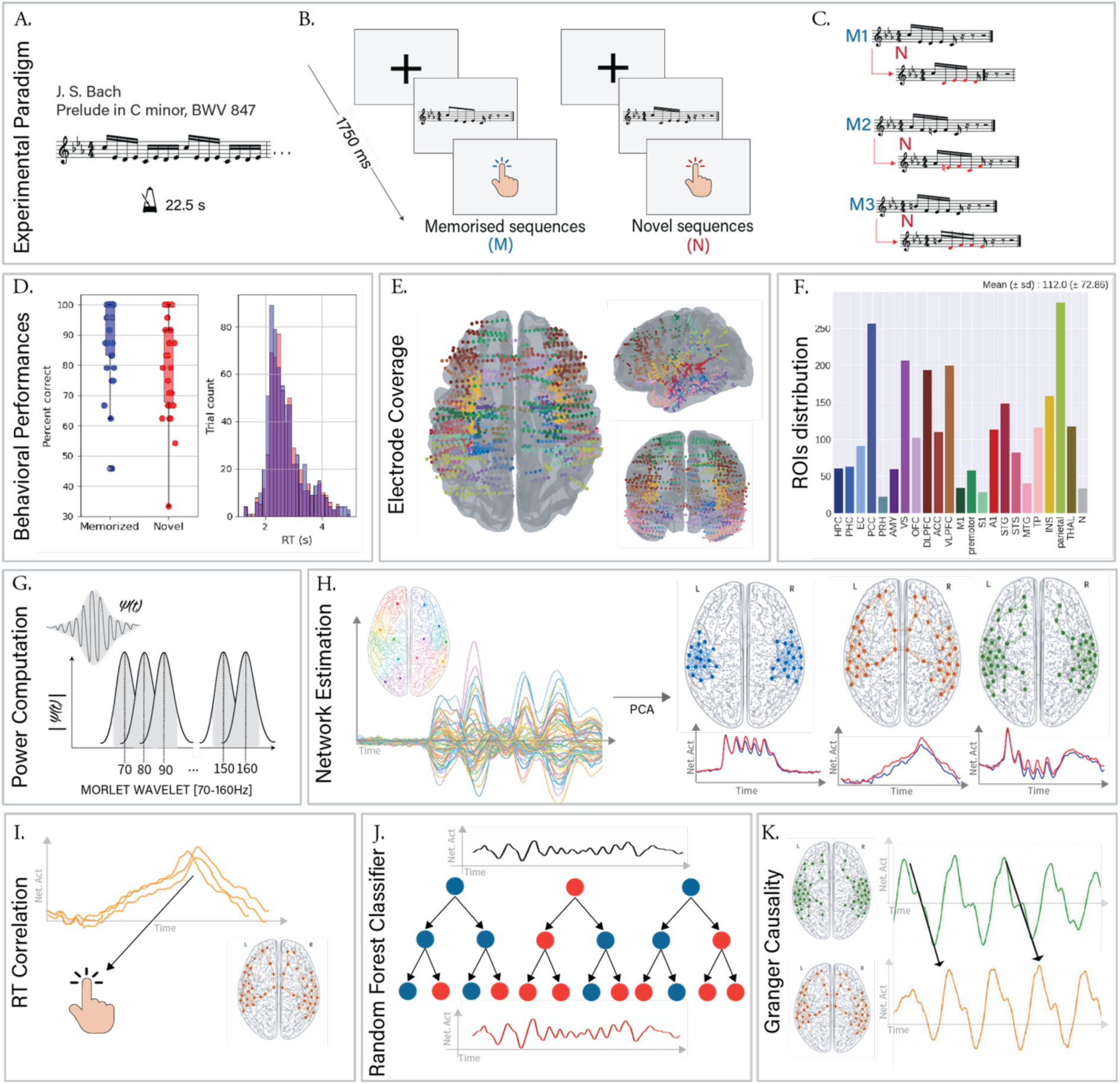
Experimental design, stimuli, and analysis pipeline a –. Participants listened to the first 4 bars of J. S. Bach’s Prelude in C minor, BWV 847, receiving the instruction to memorize them. **b** – One at a time, five-tone auditory sequences were presented in randomized order, and participants were instructed to respond with button presses whether they were ‘old’ (memorized musical sequences, ‘M’) or ‘new’ (novel musical sequences, ‘N’). **c** – Two types of auditory sequences (M, N) were used in the study. The N sequences were created through systematic variations of the M sequences. This procedure consisted of changing every musical tone of the sequence after the first (N), as illustrated by the red musical tones. **d** – Behavioral performances, percent correct trials per condition and trial reaction time distribution, for the two conditions and across participants. **e, f –** The electrodes from all the participants were grouped to conduct group-level analysis**. g** – After sEEG data pre-processing, high gamma (70-160Hz) power was extracted using Morlet wavelet decomposition. **h –** Principal component analysis was used to reduce the channel dimension into 3 coordinated patterns of activity explaining 28.4%, 8.7%, and 3.9% of variance, respectively. The weight matrix and the time course (Network activity (Net. Act) vs Time) are retained for analysis. **i –** Reaction Time (RT) correlation analysis: Pearson linear correlation between the reaction time and the dynamics of the networks. **j** - A Random Forest classifier was applied in a cross-validated decoding framework to discriminate between memorized and novel conditions based on the temporal dynamics of each network, and feature importance scores were used to identify the most informative time window. **k –** The interactions between each network are further characterized using Granger causality (GC) at the trial level to assess directed interactions between components.

### 2.2. Three functionally distinct brain networks support recognition memory

For group-level analysis, data from all participants and electrodes were concatenated along the electrode dimension in a “*pseudo-subject*” approach as follows. Whole-brain high-gamma activity (70–160 Hz) was averaged across trials for each condition separately, concatenated across time, and dimensionality reduction was performed using principal component analysis (PCA). This analysis revealed three dominant activity patterns, explaining 28.4%, 8.7%, and 3.9% of the total variance, respectively. These components were stable across participants, with leave-one-subject-out analyses yielding reproducible time series (mean spearman’s r =1 for each component) and stable to trial removal, with a leave-one-trial-out PCA weights Spearman’s correlation coefficient of 1 for each component. PCA was also performed on subject-level data. Among the first three resulting components, 89.9%, 63.3%, and 20.0% of the subject-level component time series showed a Spearman correlation coefficient greater than 0.5 with the group-level component time series (components 1, 2, and 3, respectively) (Fig S3), confirming the reliability of the “*pseudo-subject*” approach and group-level components. PCA was also performed by computing the mean over the 2 conditions instead of the concatenation, yielding a Spearman correlation between the PCA weights matrix of 1 for the 3 components (Fig S1).

PCA extracts orthogonal components that capture dominant patterns of linear covariance^28^. Consequently, the identified components are interpreted as modes of coordinated task-related activity that we can interpret as a functional brain network. The three principal components revealed dissociable large-scale functional networks with distinct spatial organizations and temporal profiles. The spatial distribution of PCA weights identified the brain regions contributing to each network (Fig. 2b,c). The sign of each PCA weight indicated whether a region contributed positively or negatively to the corresponding component, relative to the polarity of its time course. Thus, a negative contribution indicates activation of the corresponding area when the time course is negative, whereas a positive contribution indicates activation when the time course is positive. To summarize regional contributions, we computed the signed mean PCA weight within each region of interest. The temporal profiles of the components revealed their engagement throughout the unfolding auditory sequence (Fig. 2a). Together, these patterns delineated three complementary computational systems involved in sensory processing, decision-related processes, and sensory-memory integration.

**Figure 2.**
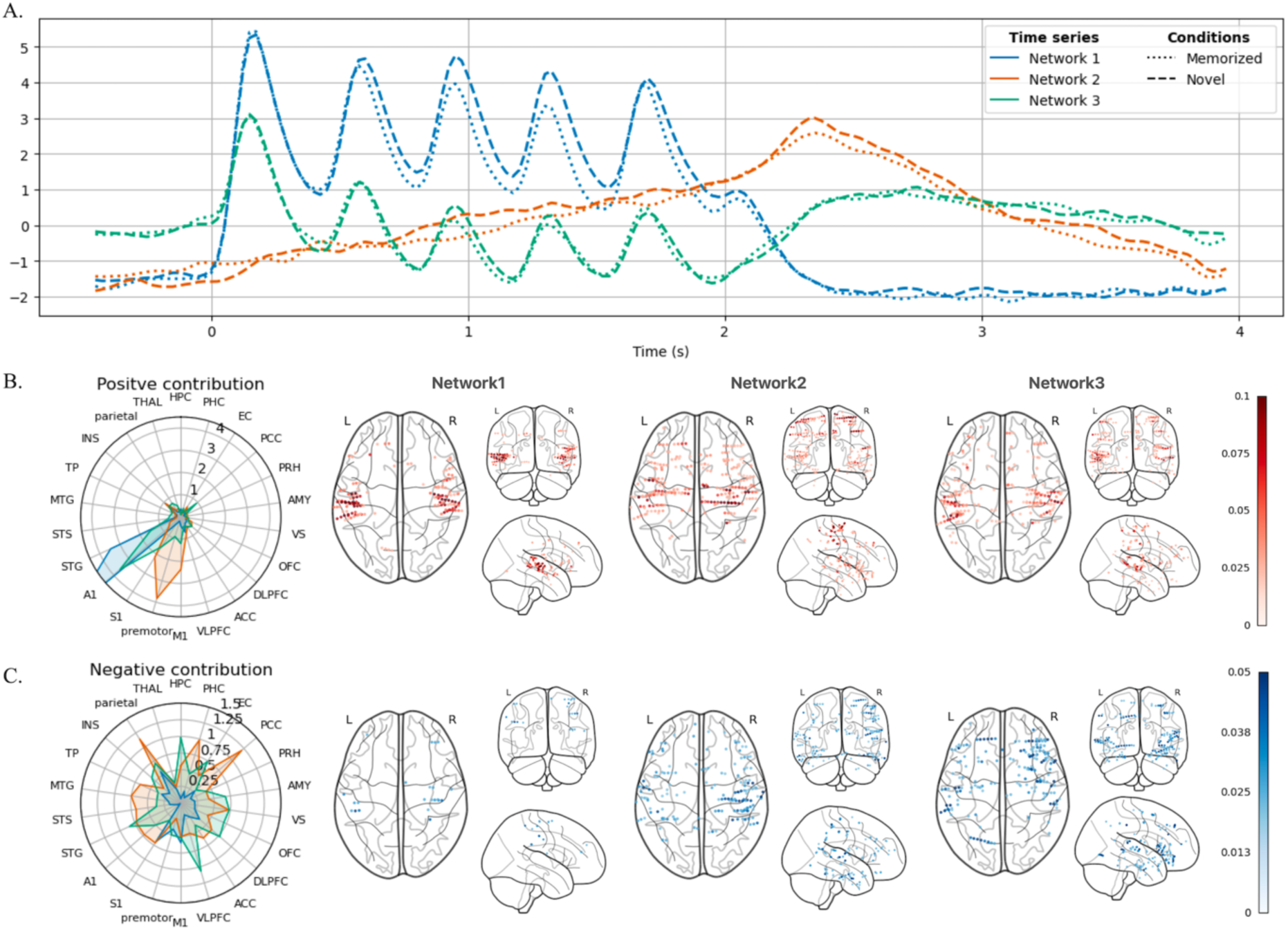
Network dynamics and spatial organization. **a** – Temporal dynamics of the three principal high-gamma networks extracted using PCA. Solid lines represent the memorized condition, whereas dashed lines represent the novel condition. Network time courses reveal distinct patterns of activity across the duration of the auditory sequence. **b, c –** Positive and negative spatial contributions of each network, respectively. Left panels show the relative contribution of anatomical regions to each component, summarized across regions of interest. Right panels depict the cortical distribution of the main electrode weights (threshold using mean + std) projected onto individual electrode locations, with warmer colors indicating stronger positive contributions and cooler colors indicating stronger negative contributions.

#### 2.2.1. Sensory processing network

The first network was focal and primarily composed of auditory regions, including the primary auditory cortex (A1) and superior temporal gyrus (STG), with a smaller but notable contribution from the insula (INS). These regions contributed positively to the component, with no substantial negative contributions. The corresponding time course exhibited clear evoked responses following each sound onset, with peak latencies computed from the trial-averaged signal at 0.18 s, 0.60 s, 0.98 s, 1.33 s, and 1.70 s for memorized sequences (M), and at 0.20 s, 0.63 s, 0.93 s, 1.35 s, and 1.73 s for novel sequences (N). This network captured online processing of auditory information.

#### 2.2.2. Decision-making network

The second network exhibited a broader spatial organization. The largest positive weights were localized in sensorimotor regions, including the primary motor cortex (M1), premotor cortex, primary somatosensory cortex (S1), and the insula (INS). Smaller negative contributions originated from the parahippocampal cortex (PHC), the parietal cortex, and the posterior cingulate cortex (PCC). Additionally, we note a small and broadly distributed contribution of auditory cortices.

The associated time course displayed a gradual increase after stimulus onset, changed polarity around 1s, and reached a maximum at 2.35 s. This pattern suggests an initial predominance of negatively weighted regions followed by progressive recruitment of positively weighted sensorimotor pathways. Accordingly, this network may reflect the transition from information integration toward decision formation and motor commitment.

#### 2.2.3 Integrative network

The third network reflected interactions between externally driven sensory representations and internally guided mnemonic-executive processes. Positive contributions were concentrated in auditory and sensory cortices (A1, STG, S1), indexing representation of incoming sensory information. Negative contributions involved MTL regions such as the hippocampus (HPC), entorhinal cortex (EC), and amygdala (AMY), together with executive regions including ventrolateral prefrontal cortex (VLPFC), dorsolateral prefrontal cortex (DLPFC) and orbitofrontal cortex (OFC), and a small contribution of the ventral visual stream (VS). We note a contribution of STG in the negative weights as well. Anatomically, this broad negative contribution is consistent with contextual and goal-directed memory retrieval.

The corresponding time course displayed five distinct peaks associated with the five tones of the stimulus sequence. The signal was initially positive, reaching its first maximum around 0.17 s, before becoming negative after the first peak, indicating a transition from dominant activation of positively weighted sensory regions toward engagement of negatively weighted memory and executive regions. The signal subsequently oscillated throughout stimulus presentation before gradually increasing to reach a late relative maximum.

The dynamics of this network suggest a push–pull interaction between incoming auditory information and internally guided memory retrieval. Each pitch elicited an initial sensory response followed by a response associated with internally driven activity. Therefore, we do not interpret this component as a purely mnemonic network, but rather as reflecting the integration of sensory and memory processes. Because it involved not only MTL regions but also executive and limbic systems, this network likely reflects the contextual integration of incoming sensory information with stored representations.

### 2.3. Auditory and decision networks discriminate memorized from novel sequences

To identify when each network carried information distinguishing memorized and novel sequences, we performed decoding using network time courses as input features for a random forest classifier within a cross-validation framework (Fig. 3). Classification performance quantified the amount of condition-specific information contained within each network, whereas feature importance scores reflected the contribution of each time point to the classifier after model training.

**Figure 3.**
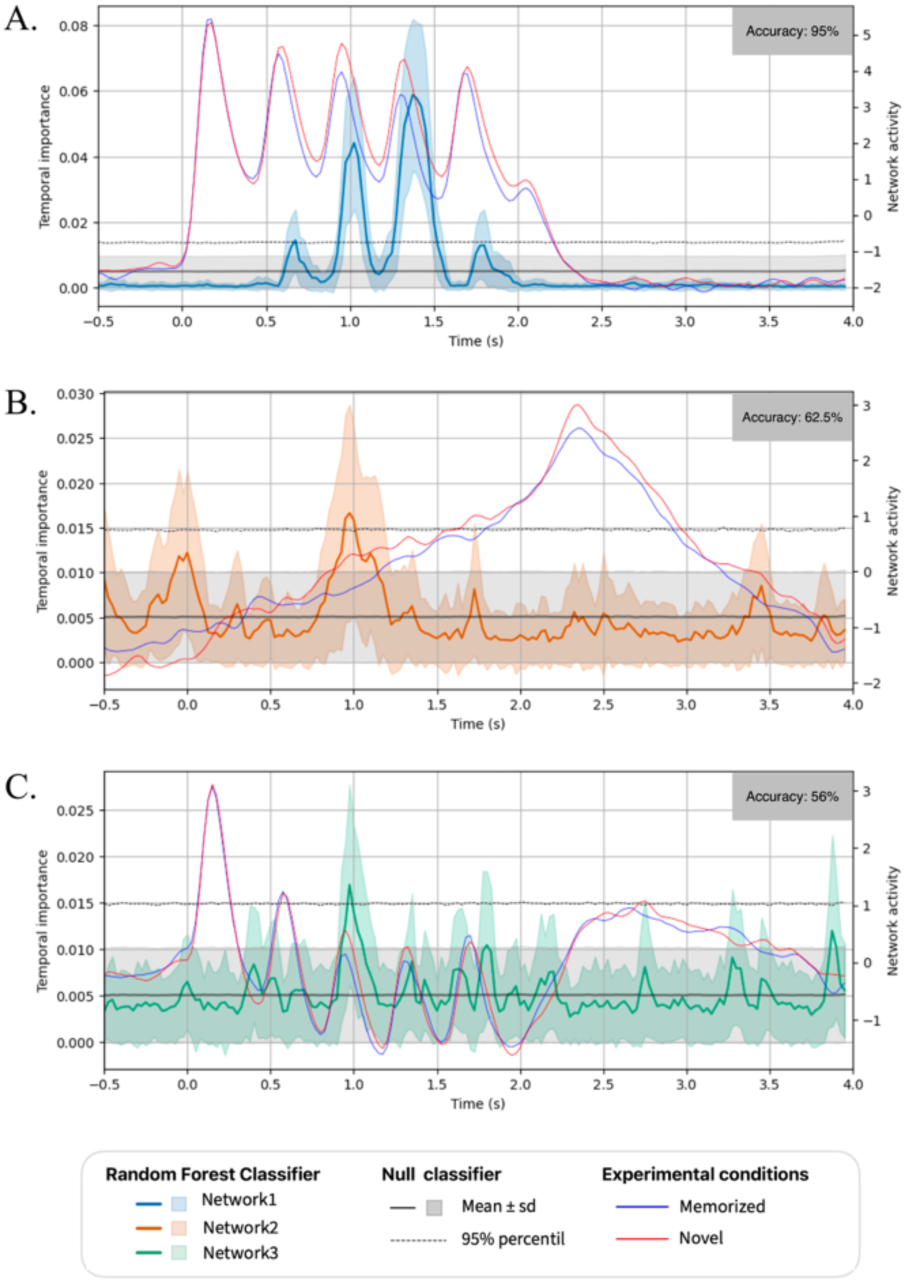
Condition discrimination among networks using Random Forest classifier. Temporal feature importance profiles obtained from the Random Forest classifier for each network. Feature importance was estimated using the impurity-based importance score and reflects the relative contribution of each time point to the discrimination between memorized and novel conditions. Distinct networks showed different temporal windows carrying discriminative information. Auditory processing, decision-making, and integrative networks, respectively, a, b, c.

#### 2.3.1. Predictive suppression in the sensory processing network during memorized sequences

Decoding of the first network achieved a mean accuracy of 96% (null-model accuracy: 50%), with an F1 score of 96% across 100 cross-validation runs. The mean accuracy over the subsampling is significant (Monte Carlo test with 1000 permutations, p=0.00099; p<0.001) and stable (SD=0.14, see Fig S5 for distribution). The most informative time points were located shortly after the peaks associated with the third and fourth tones, where feature importance exceeded the 95th percentile of the null distribution. Additional informative time points were observed following the second and fifth tones, with feature importance exceeding the null mean plus one standard deviation and slightly above the 95th percentile for the second tone. Overall, discrimination between conditions was primarily driven by amplitude differences during the descending phase of the evoked response, whereas the ascending phase was largely preserved across conditions. Compared with the novel condition, the memorized condition showed a modest reduction in amplitude at the second peak and a more pronounced attenuation at the third and fourth peaks. These findings indicate that memorized sequences were associated with reduced sensory responses, consistent with predictive modulation of auditory processing. The timing of condition discrimination suggests that differences emerged progressively during sequence processing, which may reflect early sensory and later memory-related comparisons. The second pitch already showed a difference in amplitude, suggesting an early divergence in sensory processing. The third pitch represented the earliest robust point of discrimination, occurring after sufficient sequence information had accumulated to allow comparison with stored representations.

#### 2.3.2. The decision-making network captures the timing of deviance detection

The second network yielded a mean decoding accuracy of 62.5% (null model accuracy: 50%), with an F1 score of 61% across 100 cross-validation runs. The mean accuracy over the subsampling is significant (Monte Carlo permutation test with 1000 permutations, p=0.0039; p<0.005) and relatively unstable (SD=0.33) (see Fig S5 for distribution). Feature importance analysis revealed a significant contribution around 1 s, corresponding to the transition toward engagement of sensorimotor regions. This time point exceeded the 95th percentile of the null distribution. The transition occurred earlier in the novel condition than in the memorized condition, suggesting a faster recruitment of sensorimotor-related processes during novel sequence evaluation. For novel sequences, decisions can be made after the second sound, which is always a deviant, and may be driven by its surprisingness. For memorized sequences, the decision formally requires hearing the final pitch, as the sequence must be evaluated in its entirety (although, across trials, learning may allow decisions to occur at this point as well). This point captures the moment where recognition decisions are made and evolve toward motor commitment.

#### 2.3.3. The integrative network alternates between internal cognition and perception

Decoding based on the third network yielded performance only slightly above chance level, with a mean accuracy of 56% (null model accuracy: 50%) and an F1 score of 53% across 100 cross-validation runs. The mean accuracy is not significant (Monte Carlo permutation test with 1000 permutations, p=0.051) and relatively unstable (SD=0.34, see Fig S5 for distribution). Feature importance analysis suggested a potential difference around 1 s, corresponding to the onset of the third pitch, with larger amplitudes in the novel condition compared with the memorized condition, implying an enhanced activity in the auditory cortex. However, this effect was not consistent across trials to allow reliable discrimination between conditions, with decoding accuracy just above-threshold *p*-value. This indicates that this network might be involved in sensory-mnemonic integration regardless of the novelty of the stimuli.

### 2.4 Network dynamics predict the timing of behavior

We investigated the relationship of brain dynamics with the timing of participants’ responses to define the behavioral relevance of the identified networks. The results reported are significant at α = 0.05, computed with empirical *p*-values obtained from 1,000 RT permutations and corrected for multiple comparisons using false discovery rate (FDR).

Consistent with the online processing of Network 1, the timing of sensory processing deactivation, characterized using the transition from positive to negative values (zero-crossing) (Fig. 4c left), correlated with participants’ reaction times (RTs): median r = 0.635 for memorized (M) and 0.591 for novel (N) sequences (Fig. 4d). This relationship indicates that the end of sensory processing correlates with the initiation of behavior in both conditions.

**Figure 4.**
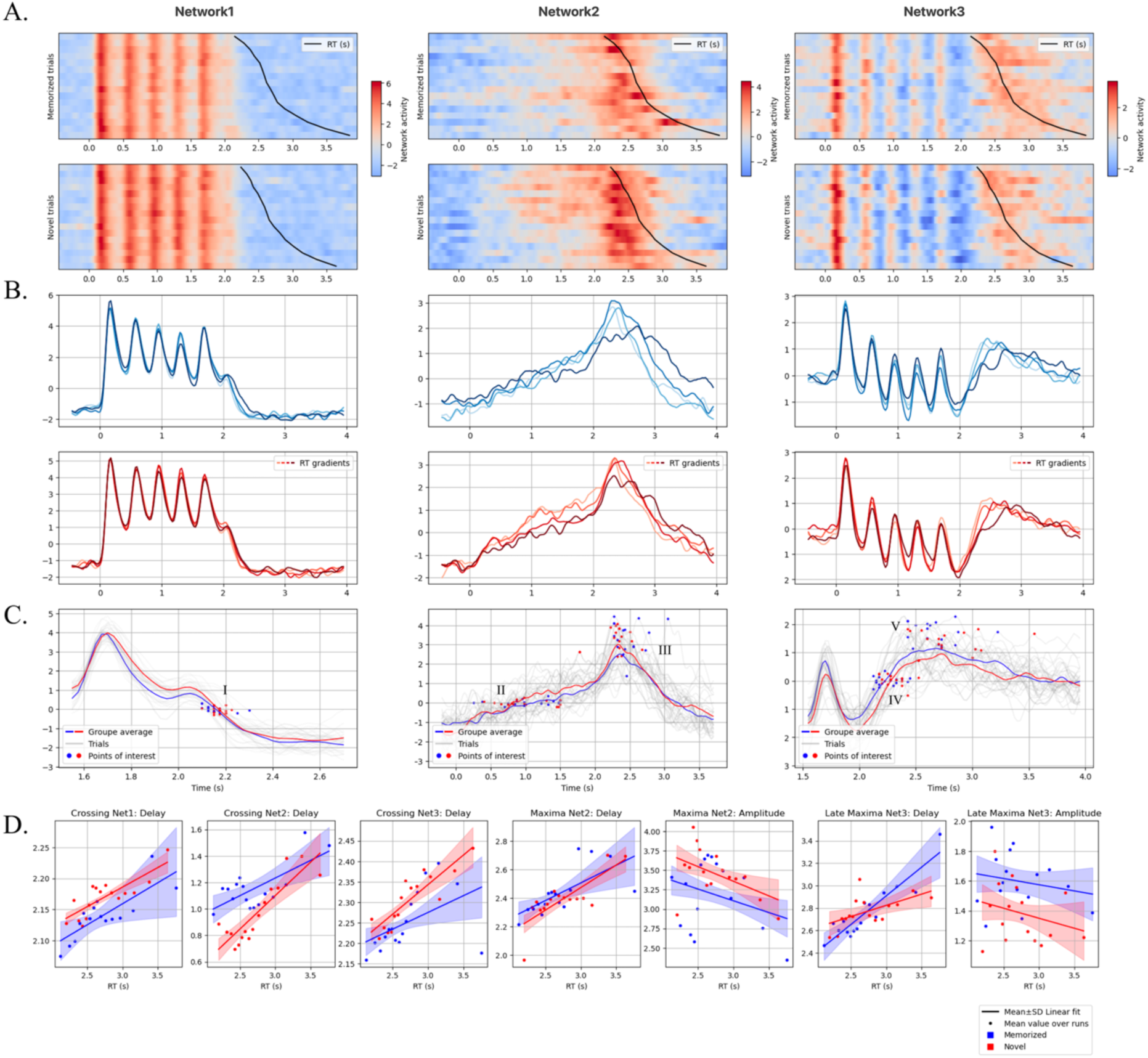
The timing of behavior correlates with the brain network dynamics. **a** – Sixteen “pseudo-subject” trials ranked by reaction time (RT) and associated with the corresponding group-level RT. **b** – Trial sub-averages. Across the 500 runs, the trials transformed by PCA are retained, ordered according to group-level RT, and averaged into four groups representing RT gradients within each condition (c: Memorized; d: Novel). **c** – To capture potential changes of the same motif, we estimated the timing and amplitude of the motif across trials. The characteristic motifs extracted were Network 1: (I) zero-crossing, marking the end of auditory cortex activity. Network 2: (II) zero-crossing, marking the transition from the contextual memory and integration areas to the sensorimotor pathway;(III) delay and amplitude of the minima, marking the end of information accumulation leading to motor commitment. Network 3: (IV) zero-crossing, marking the end of engagement of the memory-related areas; (V) delay and amplitude of the late maxima, corresponding to the final activation of the auditory cortex at the end of the process. **d** – Pearson correlation between motif features and reaction time average across runs. Each run corresponds to a randomly sampled set of trials used to build the “pseudo-subject”. The mean RT across subject trials is used to assign an RT to each resulting “pseudo-subject” trial. The correlation is computed on each set of random sampling of trials (100 iterations) and averaged for visualization. For statistical evaluation, the median across the random sampling is retained and compared against medians from RT random permutations (Fig. S5).

For decision-related activity in Network 2, significant correlations were observed, stronger for novel sequences. In Novel sequences, faster responses were associated with an earlier transition toward sensorimotor engagement (median r = 0.647) and an earlier peak of activity in the sensorimotor area (median r = 0.667) (Fig. 4C middle and 4d). In contrast, in the memorized condition, the timing of the transition toward sensorimotor engagement (median r = 0.420) and the timing of the maxima (median r = 0.354) with RT (Fig. 4c middle) were less robust. In terms of amplitude of the maxima, both conditions show significant correlation; nevertheless, those relations are weak (median r =-0.149 (M),-0.264 (N)), reflecting the importance of timing rather than amplitude.

The internal end of processing was characterized by a release of activity in the region with negative weights in Network 3. The timing of this transition (Fig 4c right) was associated with response times in the novel condition (median *r* = 0.639), but less so in the memorized condition (median *r* = 0.285). Conversely, the timing of the late increase in activity in the regions with positive weights (Fig. 4c right) in activity showed a stronger association with response times in the memorized condition (median *r* = 0.699) than in the novel condition (median *r* = 0.288; Fig. 4d). In the novel condition, response timing was more linked to the disengagement of internal information processing (negative weights), whereas in the memorized condition, it was more associated with the reactivation of sensory regions (positive weights). The weaker association with the end of internal processing in the memorized condition is consistent with the possibility that recognition had already occurred earlier in the sequence, making the termination of memory-related processing less closely tied to the behavioral response. Notably, signal amplitude showed little association with response times in either condition, emphasizing the behavioral relevance of the timing rather than the magnitude of network activity.

Together, these findings suggest that the temporal organization of network activity reflects condition-specific computational processes whose dynamics are closely associated with behavioral performance.

### 2.5. Structured network interactions underlie recognition memory

The three identified networks can be viewed as functionally specialized computational units, each supporting distinct cognitive processes required for task performance. Each network implements a specific computation from end to end. For example, Network 1 serves as a sensory processing network but does not necessarily transmit sensory information directly to Networks 3 and 2. Instead, the memory and decision-making networks encompass transformations from sensory input to memory retrieval and motor commitment, as both are connected to sensory units. Despite this specialization, these parallel computations were predicted to interact dynamically during task execution, as each computation’s state and output must influence the dynamics of the others. To characterize these interactions, we applied sliding-window Granger causality (GC) analysis to the network time courses (Fig. 5). By quantifying temporal predictive dependencies between networks, this approach identified when activity in one network precedes and predicts activity in another, providing insight into their online coordination. For each window, model lag parameters were optimized using the Bayesian Information Criterion (BIC) to ensure appropriate time lag selection. Statistical significance was assessed using surrogate-based null distributions generated via multiple-time permutation approaches, controlling for spurious dependencies while preserving relevant temporal properties. The Net GC is computed using the difference between the log-variance in both directions. GC for each direction, as well as using the reverse time sequence, is reported in the supplementary material.

**Figure 5.**
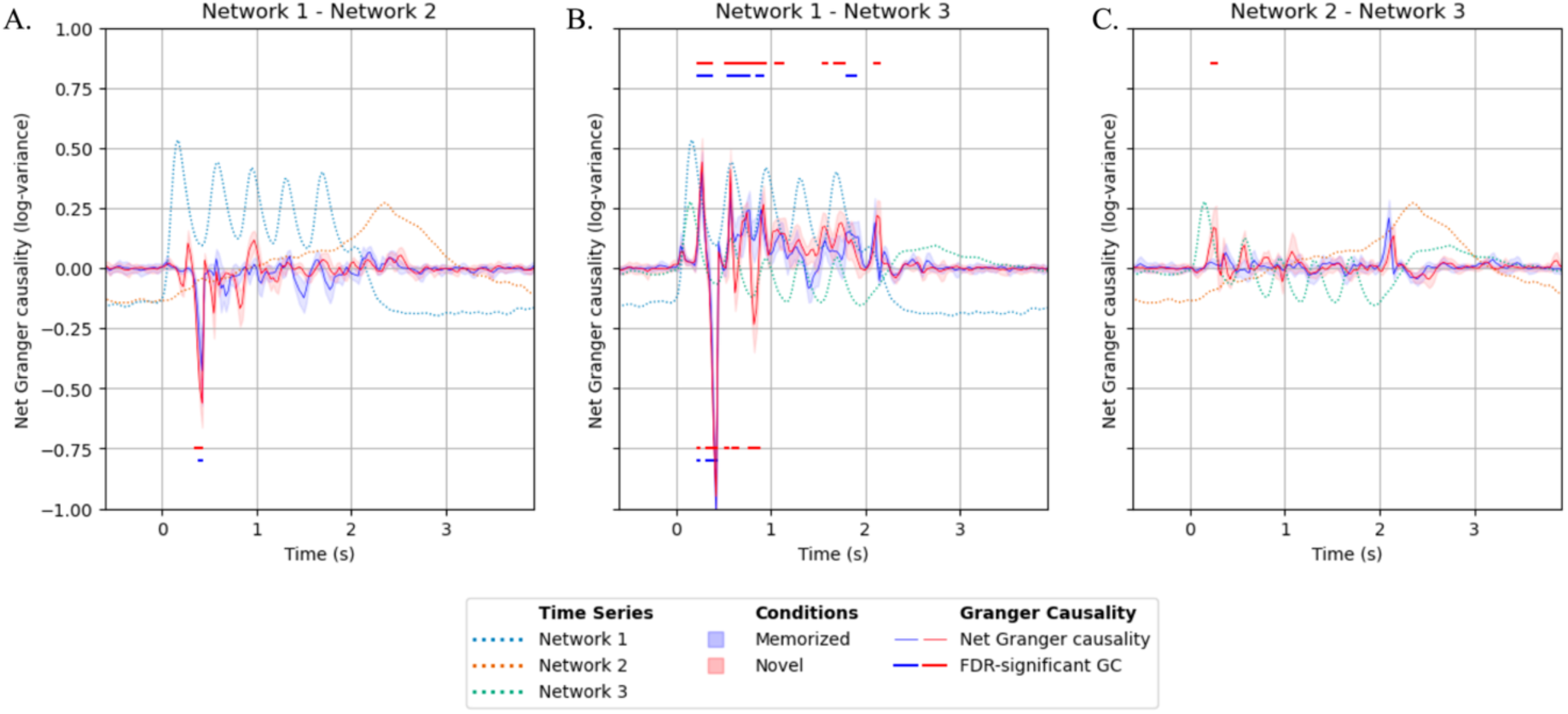
Interaction between networks within the memorized and novel conditions. Pairwise net Granger causality (GC) between **a –** sensory-processing and decision-making networks, **b –** sensory-processing and integrative networks, and **c –** decision-making and integrative networks. Directed interactions were quantified using net GC, defined as the difference between GC estimates in the two opposing directions. Positive and negative net GC values indicate predominant information transfer in opposite directions between each network pair. Significant directional interactions are indicated by lines above or below the zero net-GC line, according to the direction of information transfer. Statistical significance was determined using surrogate-based empirical p-values, followed by Benjamini–Hochberg false discovery rate (FDR) correction for multiple comparisons across temporal windows (FDR-adjusted p < 0.05). Only significant interactions with an absolute net GC magnitude exceeding 0.2 are displayed.

Significant directed interactions from auditory processing toward the integrative network emerged from the first tone and between the second and third evoked responses, with stronger interactions during the descending phase of the first response and around the peak of the second response (Fig 5B). Conversely, the integrative network exerted predictive influence on auditory processing around the minima following the first and second peaks. This feedback interaction reached its maximum within the temporal window [0.10–0.475 s]. At the end of the sequence, a final network1-to-network3 interaction was observed, potentially reflecting confirmation of the completed sequence. Time-reversed GC analyses ruled out signal-to-noise directionality confounds^29^ (Fig. S4).

The integrative network showed predictive influences on sensory processing, particularly during novel sequences, when incoming information deviated from stored expectations and required updating of internal representations. However, top-down interactions, while dominant, were not the only communication pathway observed. The bidirectional exchange between auditory processing and integrative networks (Fig 5B) further supports recognition as a continuous updating process, in which bottom-up sensory evidence and top-down expectations interact to support pattern comparison and completion throughout the sequence.

## 3. Discussion

We investigated the neural dynamics underlying recognition of temporally structured auditory sequences using high-frequency activity recorded during a memorized/novel memory paradigm. Participants were presented with five-tone melodic sequences that were either previously learned or newly generated from the variation of the 4th-to-last pitches. We reveal three large-scale functional networks supporting sensory processing, sensory-memory integration, and decision formation. These networks showed distinct temporal dynamics, condition-dependent differences, as well as directed interactions, indicating a distributed and dynamically reconfiguring architecture for ongoing and contextual recognition.

### 3.1. Recognition emerges from three interacting functional systems

A central finding is that high-frequency activity organizes into three distinct large-scale networks corresponding to three major computational demands of recognition: (1) sensory analysis of incoming auditory information, (2) goal-directed and in-context comparison of sensory input with mnemonic representations, and (3) the formation of a decision and a behavioral response^30^. Despite their interactions, these networks were independent, suggesting that recognition memory relies on multiple parallel computational systems rather than a single stepwise distributed process. Each network exhibited a distinct temporal profile and anatomical organization, consistent with a specialized functional role.

Our findings further highlight the value of adopting a network-based perspective on cognition^31^. Traditional region-of-interest approaches have been successful in identifying brain areas associated with sensory processing, memory retrieval, and decision-making^6,10,32–34^. However, our results reveal that cognition emerges from continuous interactions among distributed neural populations rather than from isolated regions acting independently^28,31,35^.

### 3.2. Continuous interaction between sensation, memory and action

#### 3.2.1. Early sensory processing is rapidly shaped by contextual predictions

Recognition begins with stimulus-locked auditory responses within A1, STG and the insula, regions consistently implicated in auditory and sequential musical processing^20,36–38^. Although Network 1 primarily reflects early sensory encoding, its activity was modulated by stimulus context. Differences between memorized and novel sequences emerged as early as the second pitch, suggesting that early auditory processing is already influenced by learned structural regularities^39^. Nevertheless, although melodies were designed without explicit violations of musical syntax^40^, they still introduce unusual features that can challenge not only learned musical regularities but also basic sensory adaptation mechanisms. As the sequence unfolded, memorized melodies showed progressive attenuation, whereas novel melodies maintained elevated responses, consistent with sustained monitoring of unexpected sensory evidence^22,41^ and dampened responses to predicted or expected events^39^. These findings suggest that the early auditory cortex does not simply encode incoming sounds but continuously integrates bottom-up sensory input with contextual expectations, providing the initial substrate for subsequent mnemonic evaluation. Importantly, the return of Network 1 activity toward baseline was correlated with reaction time in both conditions, indicating that the end of auditory activity is behaviorally relevant for recognition.

#### 3.2.2. Recognition emerges through recurrent sensory-memory interactions

In parallel, Network 3 revealed alternating dominance between externally driven sensory activity and internally driven mnemonic processing. Rather than reflecting sequential stages, these dynamics suggest continuous reciprocal interactions in which incoming sensory evidence updates internal representations while retrieved information simultaneously shapes ongoing perception^3–5,8,13,15,42,43^. Such recurrent dynamics are consistent with models that propose bidirectional communication between cortical sensory systems and medial temporal lobe structures during recognition, with the balance between bottom-up and top-down processing shifting according to computational demands^10,34^. Importantly, the observed dynamics suggest that recognition cannot be reduced to hippocampal retrieval alone but instead depends on a distributed system integrating sensory evidence, contextual information, and task goals.

The coupling between Network 3 dynamics and reaction time further suggests that these interactions are behaviorally relevant and vary with stimulus familiarity. For novel sequences, reaction time was coupled to the de-engagement of the memory and judgment part of the network, suggesting that the timing of disengagement from internally driven memory processing is related to the speed of recognition when the incoming sequence does not match an established representation. In memorized sequences, the recovery of the stored information is not determinant for the speed of the reaction, supporting the idea that the memory is intentionally reinstated at the beginning of the sequence. Furthermore, for memorized sequences, reaction time was more associated with late activation of the sensory area and deactivation of the memory area. Finally, the decoding analysis did not provide strong evidence for a difference between conditions in this network. This suggests that the comparison between sensory and mnemonic information occurs regardless of the condition, highlighting its role as a general computational process underlying recognition. Nevertheless, the relatively small proportion of explained variance also limits the analysis and reflects a limitation of the dataset. Indeed, the variability in electrode coverage across participants and the pseudo-population approach do not allow uniform sampling of all cortical regions, potentially limiting the representation of shared network dynamics across participants. Furthermore, the decoding analysis primarily captures amplitude-based differences and does not necessarily reflect the full range of interactions and temporal dynamics distinguishing the two conditions, as suggested by the Granger causality and behavioral results that show differences between conditions.

#### 3.2.2. Integrated sensory and mnemonic evidence is transformed into action

Finally, Network 2 activity shifted from distributed contextual and mnemonic regions, including the PHC, PCC and parietal cortex, toward progressive activation of the sensorimotor pathway involving S1, premotor cortex and M1. This shift is particularly relevant given that the sensorimotor cortex includes a system closely connected to higher-order cognitive networks^44^, potentially supporting the translation of internally maintained goals and mnemonic information into motor actions. The observed temporal progression may therefore reflect the gradual transformation of integrated sensory and mnemonic information into action preparation. These dynamics closely resemble evidence-accumulation processes described by diffusion and accumulator models of decision-making, in which evidence gradually converges toward a response threshold^33,45–47^. Unlike classical perceptual decision-making paradigms, however, the accumulated evidence in the present task appears to derive jointly from external sensory input and retrieved mnemonic representations^30,48–53^.

In contrast, in the novel condition, reaction time was correlated with the timing of Network 2’s transition across zero, marking the shift toward sensorimotor engagement. Thus, the timing of this transition and subsequent motor commitment appears to be an important determinant of response speed for novel sequences. This relationship was weaker for memorized sequences, possibly because recognition of familiar melodies can rely more directly on established whole representations, reducing the dependency of progressive information accumulation for the action.

Reaction time was weakly related to the peak amplitude of Network 2 activity, suggesting that the magnitude of network recruitment was less important than the timing of its transition. This pattern is consistent with a threshold-like mechanism, in which the critical factor for response generation is when the system reaches a sufficient level of evidence for action rather than the overall magnitude of neural activity^46,47^.

### 3.3. Recognition as predictive inference

A key finding was the strong directed influence from the integrative network to the auditory processing network during the earliest stages of sequence processing. Rather than a predominantly feedforward flow from auditory cortex to higher-order memory systems, our results reveal an early top-down influence, indicating that sensory processing is rapidly constrained by prior knowledge. Within predictive coding frameworks, mnemonic representations provide priors that shape the interpretation of incoming sensory evidence^2–4,40^. The observed top-down influence suggests that partial sensory input rapidly reinstates a mnemonic template, biasing auditory processing toward an expected sequence structure, consistent with models of pattern completion and memory reinstatement^5,11^.

Recognition, however, did not rely on prediction alone. Auditory regions subsequently exerted reciprocal bottom-up influences on the integrative network, consistent with iterative updating of internal models as sensory evidence accumulated^42^. These reciprocal interactions persisted longer for novel sequences, suggesting that prediction errors required continued refinement before a stable representation could be established^8^.

Together, these findings support a recurrent architecture in which prediction and sensory evidence evolve through continuous reciprocal interactions rather than a strictly feedforward hierarchy.

### 3.4 Methodological considerations

The present framework demonstrates that dimensionality reduction can identify functionally meaningful population dynamics despite the substantial spatial heterogeneity inherent to multisubject iEEG recordings. Compared with previous MEG studies^18,54^, the proportion of explained variance was lower, likely reflecting the higher dimensionality of invasive recordings, the preservation of inter-individual variability, and the reduced signal mixing afforded by iEEG.

Several considerations should nevertheless be considered when interpreting the identified components. PCA extracts orthogonal components that capture dominant patterns of linear covariance, and it is not intended to recover all functional interactions or nonlinear dependencies^28^. Consequently, the identified networks should be interpreted as the principal modes of coordinated task-related activity rather than canonical anatomical boundaries, direct structural connections, or exhaustive brain systems. This perspective also helps interpret several observations from the present study. First, Network 3 showed little condition-specific modulation, suggesting that not all functional processes contributing to recognition necessarily differentiate between memorized and novel sequences. Instead, some networks may support computations common to both conditions, whereas behavioral differences emerge from their interactions with other functional systems. Similarly, the absence of effective connectivity between the integrative and decision-making networks should not be interpreted as evidence that these systems operate independently. Rather, it indicates that their dominant patterns of coordinated activity were functionally dissociable within the PCA decomposition. This interpretation is consistent with the observation that each component included auditory regions, suggesting that different neuronal populations within structures such as A1 and STG participate in distinct computations depending on task demands^31^.

More broadly, these findings emphasize that behaviorally relevant information is not only encoded within the activity of individual regions or even single functional networks with a feedforward structure but can emerge from their temporal coordination and interactions^55^. Rather than replacing region-based approaches, data-driven decomposition provides a complementary framework for identifying transient functional assemblies that support cognition across distributed brain systems^31^. The present work demonstrates that dimensionality reduction offers a powerful approach for studying distributed computations in heterogeneous human iEEG datasets.

## Conclusion

Together, these findings suggest that recognition in more naturalistic settings is not implemented by isolated memory regions or sequential processing stages. Rather, it emerges from the dynamic coordination of distributed functional networks that continuously integrate sensory evidence, mnemonic representations, and predictive signals. By revealing how these functional systems interact over time, our results provide a network-level account of recognition that bridges perception, memory, and decision-making within a unified computational framework. More broadly, they support a dynamic view of cognition in which behavior arises from the transient coordination of distributed neuronal populations, offering new insights into how the brain supports recognition under the continuous and uncertain conditions of real-world experience.

## Materials and methods

### Participants

Thirty patients undergoing presurgical epilepsy monitoring were included in the study (N = 30, 16 females; age = 37.32 ± 11.91 years, mean ± SD). Patients with chance-level or below-chance task performance, low IQ, or impaired memory capacity were excluded. All participants provided written informed consent before participation. Data were collected at multiple clinical sites across the United States and Europe. Electrode implantation and coverage were determined exclusively by clinical requirements. Across patients, 2,574 sEEG electrodes were retained for analysis. The study was approved by the institutional review boards of the participating hospitals and the University of California, Berkeley. All procedures were conducted in accordance with the Declaration of Helsinki.

### Experimental paradigm and stimuli

Participants performed a memorized/novel auditory recognition task. During an initial encoding phase, they listened twice to a short musical excerpt and were instructed to memorize it. The stimulus consisted of the first four measures of the right-hand melody from Johann Sebastian Bach’s Prelude No. 2 in C Minor, BWV 847. Each measure contained 16 tones, yielding a total of 64 tones. Individual tones lasted 350 ms, resulting in a total duration of 22,4 s. A final closing tone lasting 1 s was appended to provide a sense of musical resolution, bringing the total duration to 23.4 s. Following the encoding phase, participants were presented with 48 musical sequences consisting of five tones each (duration = 1,75 s). For each sequence, participants indicated whether it belonged to the memorized sequence or represented a variation by pressing the numeric key 1 for “memorized” sequences [M] (old) and the numeric key 2 for “novel” sequences [N], (new). Participants used their left hand to perform the task. Half of the excerpts were extracted directly from the original musical piece, whereas the remaining excerpts were novel variations. Memorized sequences (M) were derived from the first five tones of the first three measures of the original composition. Each sequence was repeated eight times, yielding 24 trials. Novel sequences (N) were generated by systematically modifying the memorized sequences. For each of the three memorized sequences, eight variations were created by altering the 4 tones following the first tone. Several transformation strategies were employed. In some cases, the melodic contour was inverted, reversing the original pitch trajectory. In others, the remaining tones were scrambled, or a single tone was repeated multiple times, occasionally with octave variations. Additional manipulations involved scrambling the interval structure between tones. In most cases, the harmonic framework of the original sequence was preserved.

### Preprocessing

The data were converted to BIDS format. We obtained electrode locations based on post-operative computerized tomography images co-registered with individual MRI scans^37^. The coordinates were translated into standard MNI space for visualization. Preprocessing was done using a custom pipeline implemented using MNE-Python^56^. We first applied bipolar re-referencing^57^ and removed channels that exhibited epileptic activity (e.g., interictal spikes or pathological slowing) or corrupted signals or were placed outside the brain or in white matter as identified through visual inspection. Further steps included resampling to 600 Hz; high-pass filtering (0.1 Hz cutoff, with no low-pass); notch filtering at 60, 120, 180, and 240 Hz for US datasets and 50, 100, and 150, and 200 for European datasets; extraction of task-related events; epoching (from-1 s to 5 s); and normalization of each channel using the mean and standard deviation across timepoints and trials. To account for bipolar re-referencing, the final location of a contact was determined as the midpoint between its original location and that of its neighbor.

Time-frequency decomposition was performed using Morlet wavelets at 38 center frequencies spanning 0.5-180 Hz, with a variable bandwidth. Power estimates were log-transformed and z-scored separately for each channel and frequency using the mean and standard deviation computed across all trials and baseline within the analyzed epoch. High-frequency activity (HFA) was defined as the average normalized power across 10 frequency bins (for 70 to 160 every 10Hz; bandwidths: 10Hz)^58^.

### Anatomical Locations and Regions of Interest

Electrode location and labelling were determined by visual inspection and corroborated by an expert neurologist. We further grouped the electrodes into the following anatomical regions.

**Table 1.**
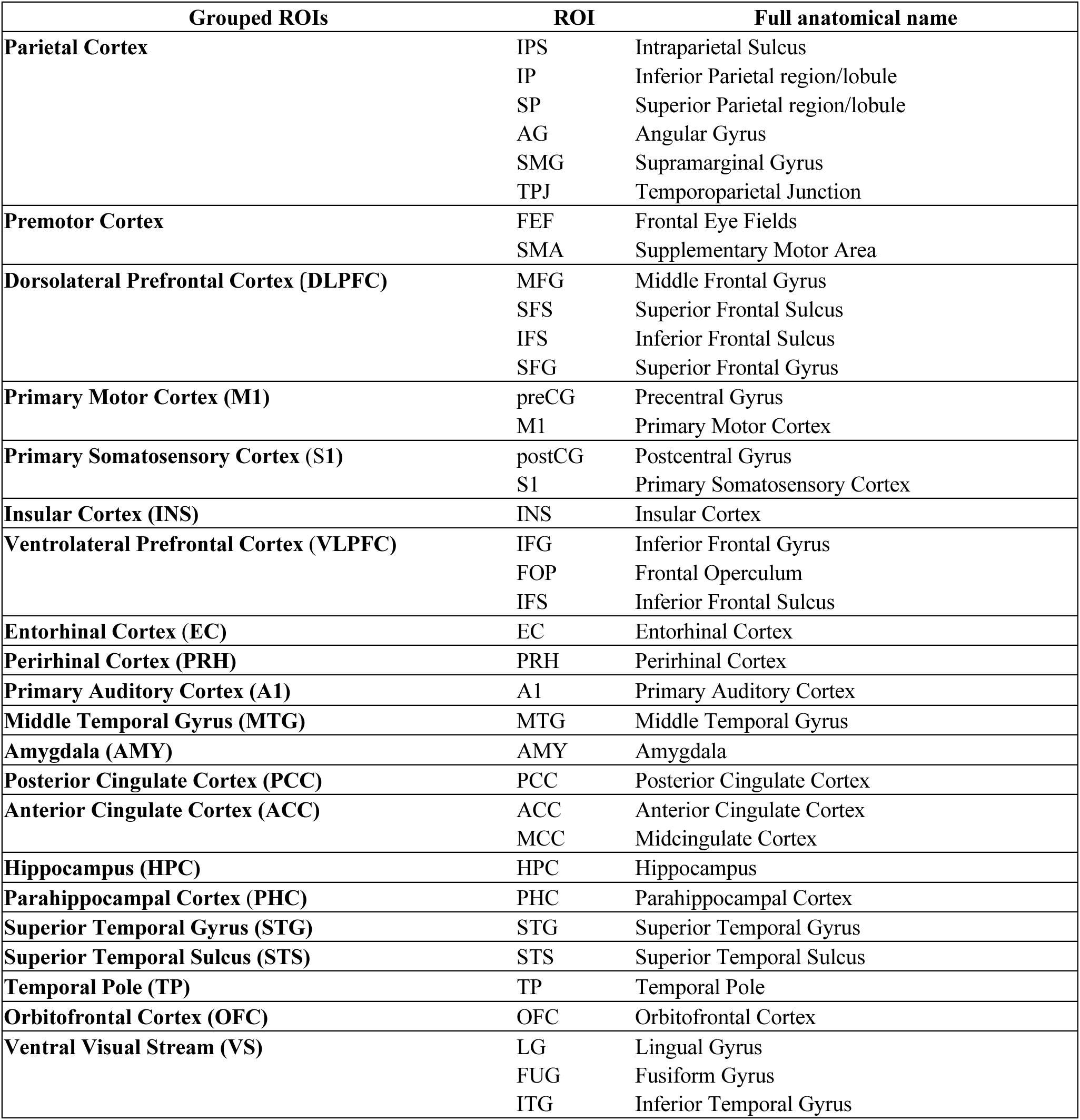
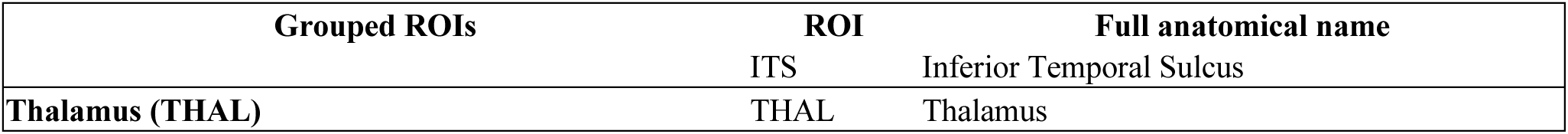
Region of interest labelling.

| **Grouped ROIs** | **ROI** | **Full anatomical name** |
|---|---|---|
| **Parietal Cortex** | IPS | Intraparietal Sulcus |
|  | IP | Inferior Parietal region/lobule |
|  | SP | Superior Parietal region/lobule |
|  | AG | Angular Gyrus |
|  | SMG | Supramarginal Gyrus |
|  | TPJ | Temporoparietal Junction |
| **Premotor Cortex** | FEF | Frontal Eye Fields |
|  | SMA | Supplementary Motor Area |
| **Dorsolateral Prefrontal Cortex (DLPFC)** | MFG | Middle Frontal Gyrus |
|  | SFS | Superior Frontal Sulcus |
|  | IFS | Inferior Frontal Sulcus |
|  | SFG | Superior Frontal Gyrus |
| **Primary Motor Cortex (M1)** | preCG | Precentral Gyrus |
|  | M1 | Primary Motor Cortex |
| **Primary Somatosensory Cortex (S1)** | postCG | Postcentral Gyrus |
|  | S1 | Primary Somatosensory Cortex |
| **Insular Cortex (INS)** | INS | Insular Cortex |
| **Ventrolateral Prefrontal Cortex (VLPFC)** | IFG | Inferior Frontal Gyrus |
|  | FOP | Frontal Operculum |
|  | IFS | Inferior Frontal Sulcus |
| **Entorhinal Cortex (EC)** | EC | Entorhinal Cortex |
| **Perirhinal Cortex (PRH)** | PRH | Perirhinal Cortex |
| **Primary Auditory Cortex (A1)** | A1 | Primary Auditory Cortex |
| **Middle Temporal Gyrus (MTG)** | MTG | Middle Temporal Gyrus |
| **Amygdala (AMY)** | AMY | Amygdala |
| **Posterior Cingulate Cortex (PCC)** | PCC | Posterior Cingulate Cortex |
| **Anterior Cingulate Cortex (ACC)** | ACC | Anterior Cingulate Cortex |
|  | MCC | Midcingulate Cortex |
| **Hippocampus (HPC)** | HPC | Hippocampus |
| **Parahippocampal Cortex (PHC)** | PHC | Parahippocampal Cortex |
| **Superior Temporal Gyrus (STG)** | STG | Superior Temporal Gyrus |
| **Superior Temporal Sulcus (STS)** | STS | Superior Temporal Sulcus |
| **Temporal Pole (TP)** | TP | Temporal Pole |
| **Orbitofrontal Cortex (OFC)** | OFC | Orbitofrontal Cortex |
| **Ventral Visual Stream (VS)** | LG | Lingual Gyrus |
|  | FUG | Fusiform Gyrus |
|  | ITG | Inferior Temporal Gyrus |
| **Grouped ROIs** | **ROI** | **Full anatomical name** |
|  | ITS | Inferior Temporal Sulcus |
| **Thalamus (THAL)** | THAL | Thalamus |

### Principal Component Analysis

Principal component analysis (PCA) was performed on trial-averaged signals at the pseudo-subject level (i.e., concatenated all subjects in the channels dimension, leading to an overall number of electrodes of 2,574). Mean signals across trials were concatenated along the temporal dimension for the 2 conditions before PCA computation. The resulting component weights were subsequently applied either to trial-averaged data or to individual trials. The same procedure was run using the average over condition to ensure the stability of the results. The results are reported in the supplementary material.

The explained variance ratio was used to guide the selection of retained components. Component stability was assessed across both trials and subjects using a leave-one-out (LOO) procedure over 100 iterations. In each iteration, either a single trial or a single subject was removed before PCA computation. Consistency was quantified using Spearman correlations between the resulting components, based on component weights for trial-level LOO analysis and time series for subject-level LOO analysis, as removing a subject doesn’t allow the alignment of weights. Each of the subjects was tested for stability.

The subject-level LOO analysis ensured that extracted components reflected patterns shared across participants rather than being driven by individual subjects. The trial-level LOO analysis verified that trial-averaged components captured stable trial-specific dynamics.

PCA was run on individual subjects (Fig. S3) following the same pipeline for conditions and trials. Spearman correlation coefficient between per-subject PCA weights and pseudo-subject PCA weights was assessed for each of the first three components. We counted how many times a correlation greater than 0.5 was found, leading to a percentage of group components appearing in subject components.

### Behavioral analysis

Behavioral performance was assessed in terms of accuracy and response time (RT). Trial-level recognition accuracy was analyzed using binomial generalized linear mixed-effects models (GLMMs), with a logit link function and a random intercept for participants to account for the non-independence of repeated observations within participants. First, overall performance was tested against chance level (50%) using an intercept-only model. The fixed-effect intercept was expressed on the log-odds scale and tested against zero using a one-sided Wald test, with a positive coefficient indicating performance above chance. Estimated accuracy was obtained by back-transforming the model intercept using the inverse logit function. The corresponding odds ratio (OR) and 95% confidence interval (CI) were calculated from the model coefficient and its standard error.

To assess differences in recognition accuracy between conditions, a second binomial GLMM included condition (Memorized vs. Novel) as a fixed effect, with Memorized trials specified as the reference category and a random intercept for participants. The condition effect was evaluated using a two-sided Wald test. The resulting OR therefore represents the odds of a correct response for Novel relative to Memorized trials, with OR values below 1 indicating lower accuracy in the Novel condition.

Reaction times (RTs) were analyzed using a linear mixed-effects model (LMM) fitted with log(RT) as the dependent variable, condition (Memorized vs. Novel) as a fixed effect, and a participant-specific random intercept. The condition effect was evaluated using a two-sided test of the fixed-effect coefficient. Mean RTs were calculated separately for each condition, and 95% CIs were obtained from the fitted model. All mixed-effects models were fitted in Python using statsmodels.

To investigate the relationship between network dynamics and behavior, trials from a subset of 25 subjects (each contributing at least 16 trials per condition) were ranked according to RT and combined across subjects to create *“pseudo-subject”* trials with a group-level distribution of reaction times (computed as the mean across subjects). Each trial was then projected onto the three network components using the PCA weights, yielding RT-sorted network time courses. Analyses were performed separately for the two experimental conditions and for each component identified by the principal component analysis (PCA).

Several features were extracted from the resulting component time courses. We selected the motifs of interest for their stability and interpretability. More sophisticated motif-based approaches were not feasible due to the substantial trial-to-trial variability in the network time courses. (**Table 2**).

**Table 2.** Motifs of interest for reaction time correlation.

| **Networks** | **Network 1** | **Network 2** |  | **Network 3** |  |
|---|---|---|---|---|---|
| **Motifs** | Late zero-crossing | Early zero-crossing | Maxima | Late zero-crossing | Late maxima |
| **Associated functional meaning** | End of activity of the sensory processing. | Transition to sensorimotor activation. | Maximum of sensorimotor engagement, delay and strength of the activity. | Final push–pull back-and-forth, reflecting the final processing stage in the comparison between the two streams of information. | Late activation of sensory regions and inactivation of the memory related regions. |

We repeated this procedure through 100 trial resampling iterations. For each resampling, Pearson’s correlation coefficient (*r*) was calculated between the group-level RT vector and each extracted neural feature. This yielded an empirical distribution of correlation coefficients across iterations. The median correlation coefficient was used as the summary statistic for each condition and neural feature. Statistical significance was assessed using a permutation procedure that preserved the neural measurements while randomly permuting the RT values within each resampling iteration. For each of the 100 iterations of random trial sampling, we ran P=1000 RT permutations and recalculated Pearson’s correlation, generating a null distribution of correlation coefficients. The median over random subsampling was extracted. Two-sided empirical *p*-values were calculated by comparing the observed median correlation with the values of the median permutation distribution.

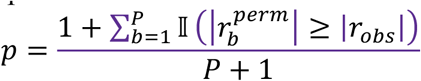

where I(⋅) is the indicator function that assigns 1 if the condition is respected. The resulting p-values were corrected for multiple comparisons using the Benjamini–Hochberg false discovery rate (FDR) procedure, with α = 0.05.

### Decoding

Because only correctly classified trials were retained, the number of available trials varied across participants. To construct a full *pseudo-subject* dataset, data augmentation was used when a participant contributed fewer than the required 24 trials per condition^59^. Two augmentation strategies were evaluated: (1) random duplication of an existing trial from the same condition and (2) replacement with the subject-level mean trial for that condition. Subject-level mean (2) was retained for its better generalization. Augmented samples represented approximately 13% of the final *pseudo-subject* dataset. PCA weights were always computed using the original, non-augmented data to avoid artificially altering the variance structure. Augmentation was restricted to training data within each cross-validation fold and was never applied to the test set.

To determine whether PCA-derived time series differed between conditions and preserve temporal structure, classifiers were trained using time points as features within each principal component. A Random Forest classifier was selected because it can model non-linear relationships and threshold-based decision boundaries while remaining relatively robust to noisy features through ensemble averaging^60^. This makes it well suited for identifying informative time points and sparse patterns of discriminative information. Hyperparameters such as the number of trees (50, 100), the max depth (None, 5, 10), and the maximum number of randomly selected features at each tree (sqrt, log2) were optimized during training. To estimate performance stability and given the limited sample size (48 trials after channel concatenation between subjects), the procedure was repeated 100 times. Each iteration employed a leave-two-out cross-validation scheme, with one trial from each condition held out for testing and the remaining trials used for training.

**Table 3.** Decoding algorithm procedure.

| Step | Procedure | Description |
|---|---|---|
| 1 | Test-set isolation | One trial per condition is isolated as the test set. |
| 2 | PCA computation | PCA is computed on the trial-averaged *pseudo-subject* training data only. |
| 3 | Data augmentation | Data augmentation is applied to the training set. |
| 4 | PCA projection | Both training and testing trials are projected onto the PCA space defined from the training data. |
| 5 | Hyperparameter optimization | Hyperparameters are optimized using a grid search during the first leave-two-out iteration. |
| 6 | Null-distribution estimation | An identical model is trained and tested using shuffled labels, with label permutation performed prior to PCA computation while preserving the same train–test split. Label permutation is repeated 50 times to estimate variability in random performance. |

Model selection was based on classification accuracy. For each time point, global classification accuracy was computed by pooling predictions across all subsampling iterations. Statistical significance was assessed using a permutation test. The observed test statistic, (*T*_obs_), was defined as the mean classification accuracy across the (R=100) subsampling iterations. For each of the (P=1000) label permutations, a null test statistic, (*T*_null,*p*_), was obtained by averaging the corresponding permuted-label classification accuracies across the 100 subsampling iterations. This resulted in a null distribution of 100 mean accuracies. A one-sided p-value testing whether classification accuracy was greater than expected under the null hypothesis was calculated as

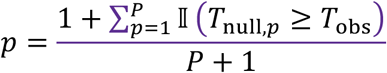

where I(⋅) is the indicator function. Thus, the p-value represents the proportion of permutation-based global accuracies that were equal to or greater than the observed global accuracy, with a +1 correction for Monte Carlo permutation testing.

For each time point, feature importance was extracted from the Random Forest model using the mean decrease in impurity and compared with a null distribution of feature importance values obtained from models trained on permuted class labels. Time points were retained when their coefficients exceeded the null-distribution mean plus one standard deviation. Statistical significance at α = 0.05 was defined as coefficients exceeding the 95th percentile of the null distribution.

### Granger Causality Analysis

Granger causality (GC) was used to quantify directed interactions between principal components derived from PCA-based reconstruction of grouped-trial time series^61,62^. Analyses were performed separately for each experimental condition using the 25 participants who contributed enough valid trials (16 per condition), thereby avoiding trial-level data augmentation. To improve the stability of the estimates, the analysis was repeated across 20 independent resampling runs. At each run, trial data were reconstructed using the predefined number of trials for each condition, and GC was estimated separately for the two trial categories (“memorized” and “novel”) and for each predefined direction between components. Time-resolved GC was computed within sliding temporal windows corresponding to the duration of a single note (350 ms). Within each window, directed GC from a source signal *X* to a target signal *Y* was estimated by comparing a restricted autoregressive model, containing only past values of *Y*,

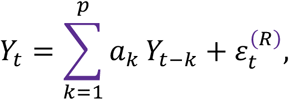

with a full model that additionally incorporated past values of *X*,

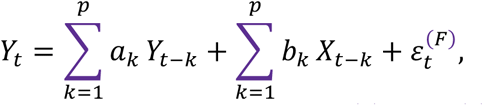

where *p* denotes the autoregressive model order and *ε_t_*^(R)^ and *ε_t_*^(*F*)^ are the residuals of the restricted and full models, respectively. Granger-causal influence from *X* to *Y* was quantified as the log-ratio of their residual variances,

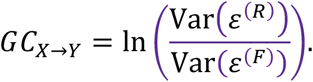

Thus, positive GC values indicate that incorporating the past of *X* reduces the prediction error for *Y*, consistent with a directed predictive influence from *X* to *Y*.

GC was estimated separately in both directions for each component pair. When directional asymmetry was summarized as net GC, it was defined as

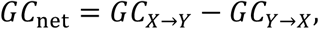

such that positive and negative values indicate a predominance of predictive influence in the *X* → *Y* and *Y* → *X*directions, respectively. Net GC therefore represents directional asymmetry rather than an independent GC estimate.

The autoregressive model order was selected independently within each temporal window. Candidate lag orders were evaluated subject to a minimum data-to-parameter constraint to reduce overparameterization. Specifically, candidate lags were retained only when

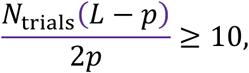

where *N*_trials_ is the number of trial observations, *L* is the window length in samples, and *p* is the candidate lag order. Among valid candidate orders, the model yielding the lowest Bayesian Information Criterion (BIC) was selected^63^. GC estimates, BIC values, F-statistics, parametric p-values, and the selected lag order were retained for each temporal window. Statistical significance was evaluated using surrogate-based empirical null distributions. For each observed GC estimate, 100 surrogate realizations were generated using the specified permutation procedure. Depending on the analysis, surrogate data were constructed using block permutation, circular temporal shifting, or temporal shuffling. For each surrogate realization *s*, the same GC estimation procedure was applied to obtain *GC*^(^*^s^*^)^, yielding a window-specific empirical null distribution,

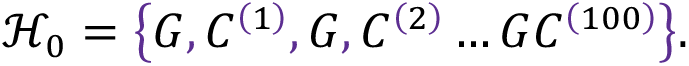

Because GC was evaluated over multiple overlapping temporal windows, the resulting empirical (p)-values were corrected for multiple comparisons across time using the Benjamini–Hochberg false discovery rate (FDR) procedure. Specifically, FDR correction was applied across the set of temporal-window (p)-values with the FDR controlled at (*α* = 0.05). Temporal windows with FDR-adjusted (p<0.05) were considered statistically significant. Surrogate analyses were performed separately for each permutation method rather than combining the three null models within a single statistical test.

To control for spurious GC associated with differences in signal-to-noise ratio^29^ (i.e., weak asymmetries), we computed GC on the time-reversed component time series before GC estimation. For a signal *X* = (*X*_1_,…,*X_T_*), the reversed series was defined as

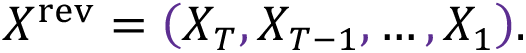

The remainder of the analysis pipeline, including windowing, autoregressive model selection, GC estimation, and surrogate generation, was then applied without modification to the time-reversed signals. A shift in the sign of reverse net GC values indicates true directional interactions and rules out signal-to-noise confounds. The complete procedure therefore yielded time-resolved observed GC estimates and corresponding surrogate null distributions for each component direction, trial category, experimental condition, resampling run, and, where applicable, time-reversal analysis.

## Supplementary materials

**Figure S1.**
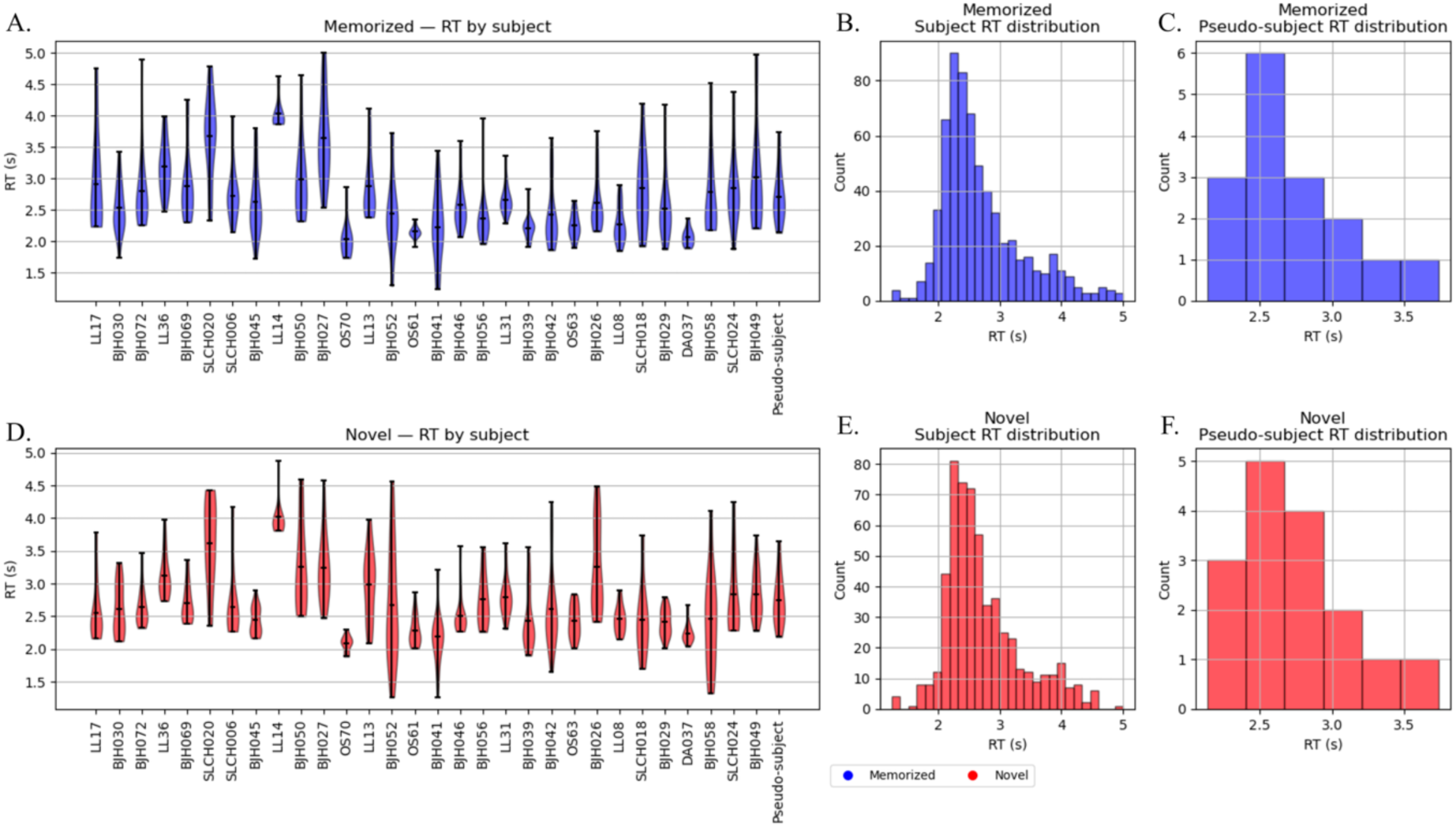
Behavioral Results. **a,d** –Reaction time per subject and at group level across trials and their distribution. A subset of subjects has been selected to ensure the same number of trials and an associated reaction time at group level. The 16 trials have been ranked by reaction time and associated in order across subjects to build the “pseudo-subject” trials. **b,e** – Reaction Time per trial count across the population; **c,f** –Reaction time at group level. The group-level reaction time is the mean over the ranked subject trial reaction time.

### Participant behavioral performances

Only successful trials have been selected. The group-level reaction time has been computed by taking the mean over the subject reaction times. Trials have been ordered by reaction time before being associated to build the *“pseudo-subject”* to preserve a distributed reaction time at group level.

### Morlet wavelet parameters and power computation

Center frequencies: 0.5, 1, 2, 3, 4, 5, 6, 7, 8, 9, 10, 11, 12, 13, 15, 17, 19, 21, 24, 27, 30, 35, 40, 45, 50, 55, 60, 70, 80, 90, 100, 110, 120, 130, 140, 150, 160, 180

Bandwidths:

- 0.5 for 0.5 Hz

- 1 for 1–13 Hz

- 2 for 15–19 Hz

- 3 for 21–30 Hz

- 5 for 35–60 Hz

- 10 for 70–180 Hz

Canonical frequency definition:

- Delta: 0.5, 1, 2, 3

- Theta: 4, 5, 6, 7

- Alpha: 8, 9, 10, 11, 12

- Low Beta: 13, 15, 17, 19

- High Beta: 21, 24, 27, 30

- Low Gamma: 35, 40, 45, 50, 55, 60

- High Gamma: 70, 80, 90, 100, 110, 120, 130, 140, 150, 160

### PCA Computation

The same pipeline has been run, computing the mean over the 2 conditions instead of the concatenation to ensure consistency for the network. The results are shown in Figure S1 and show a similar distribution in terms of components, explained variance, and temporal and spatial distribution. The mean over condition reduces noise by averaging and gives a better explained variance ratio but reduces the specific patterns of each condition as well. The explained variance ratio figure (**Fig. S1b**) shows the elbow at PC=3 for the 2 methods.

**Figure S2.**
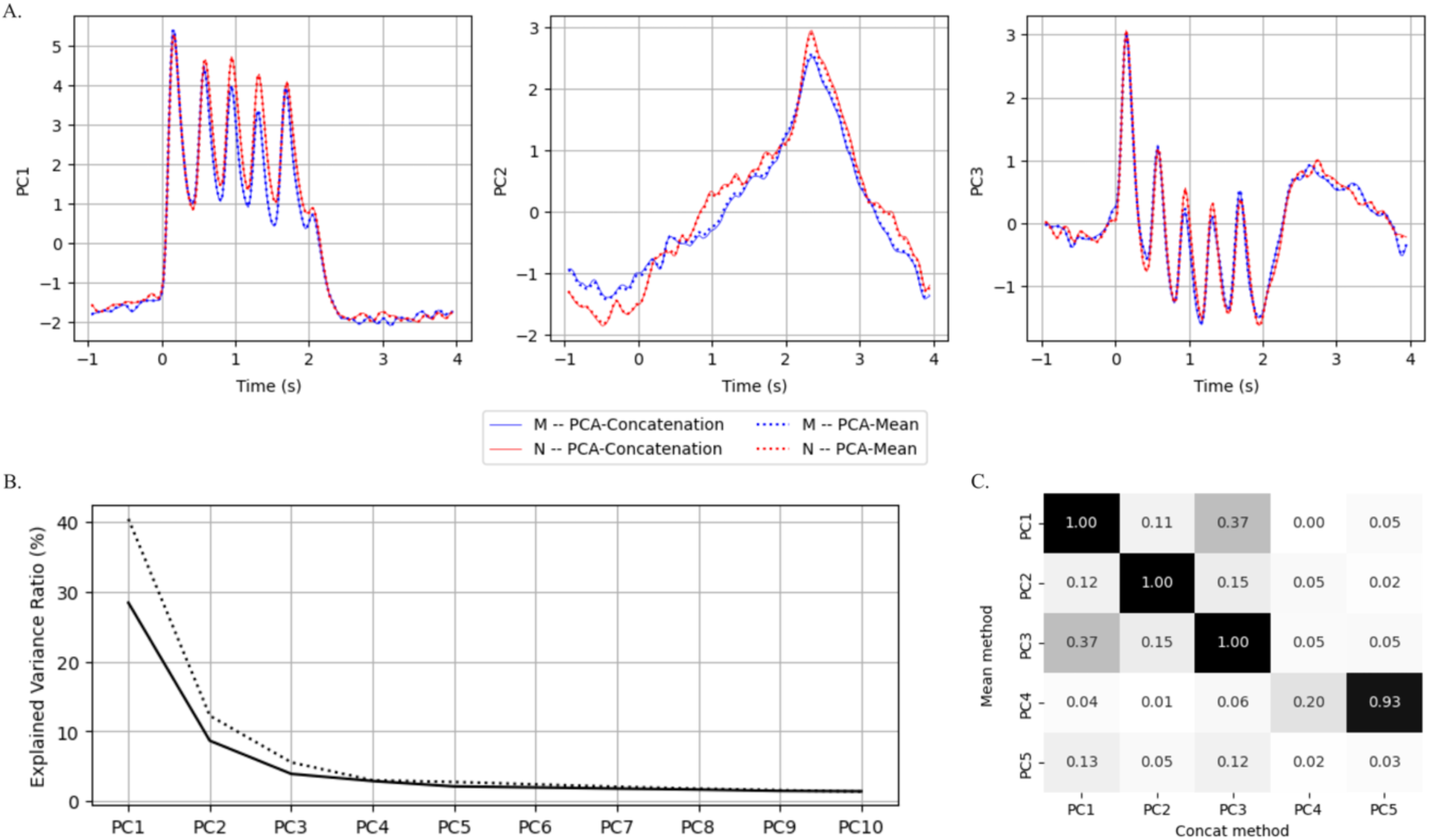
PCA stability across computation. PCA computation using the mean or the concatenation over the condition. **a** – Time courses, **b** – Explained variance ratio, **c** – Spearman r correlation on PCA weights.

### Subject-level PCA correlation

The same analysis was applied to subject-level data, and times were correlated with the group level.

**Figure S3.**
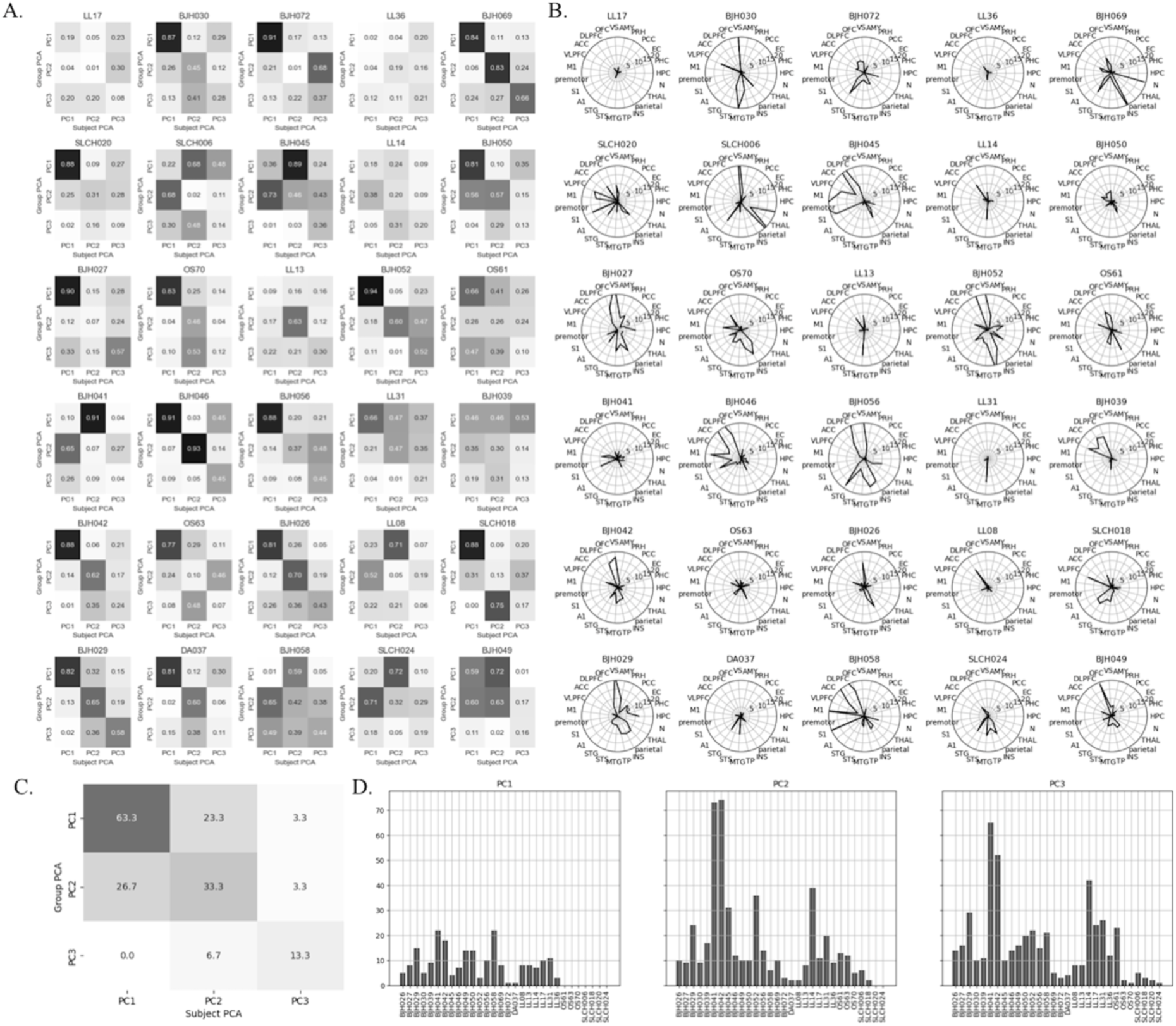
Subject-level PCA. PCA was performed separately for individual subjects using data concatenated across conditions. **a** – Spearman correlation coefficients between the subject-level component time courses and the corresponding group-level PCA component time courses. **b** – Electrode composition for each subject, showing the number of electrodes located in each region of interest. **c** – Percentage of subject-level components with an absolute Spearman correlation coefficient greater than 0.5 with the corresponding group-level component. **d** – Subject contributions to the group-level PCA weights, quantified as the number of electrodes with a significant contribution (above the mean + SD) belonging to each subject.

### Decoding

Over the cross-validation, the predictions were retained, leading to two accuracy scores. An accuracy score for each of the iterations. The accuracy distribution was used to assess the stability of the model (Fig S). A general accuracy used by pooling all the predictions together. Hyperparameters are optimized using the accuracy score on the first training split. The hyperparameters retained for the rest of the training are:

- Network 1: 100 trees, no maximal depth and sqrt randomly selected features.

- Network 3: 100 trees, no maximal depth and log2 randomly selected features.

- Network 3: 50 trees, 10 depth, and sqrt randomly selected features.

The same decoding has been performed with duplicate trials as augmentation; we obtained 95%, 69%, and 54% accuracy following the same procedure with the RandomForest Classifier. And the same profile of temporal importance emerges.

**Figure S4.**
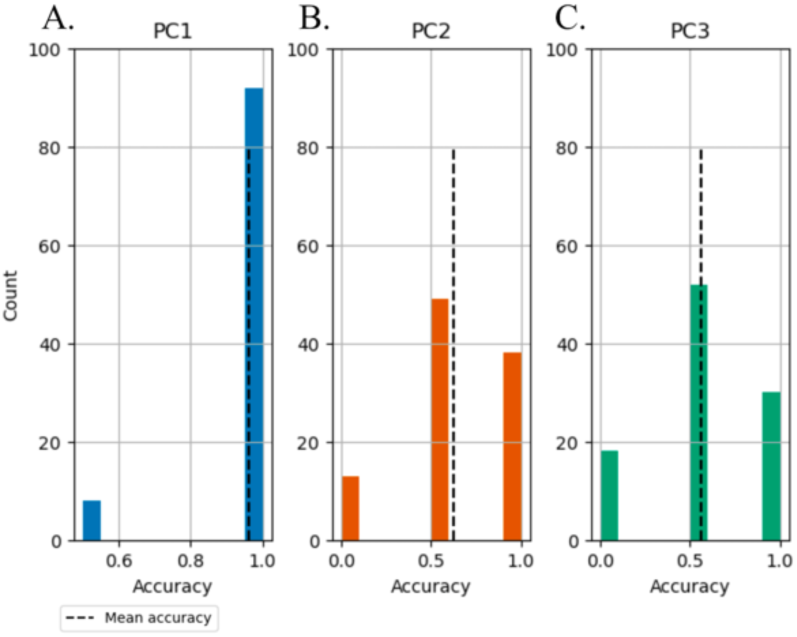
Decoding accuracy distributions over cross-validation. Accuracy distribution from the 100 cross-validation procedure. **a, b, c**, for network 1, 2, 3, respectively. The dashed line indicates the mean accuracy.

Granger Causality

**Figure S5.**
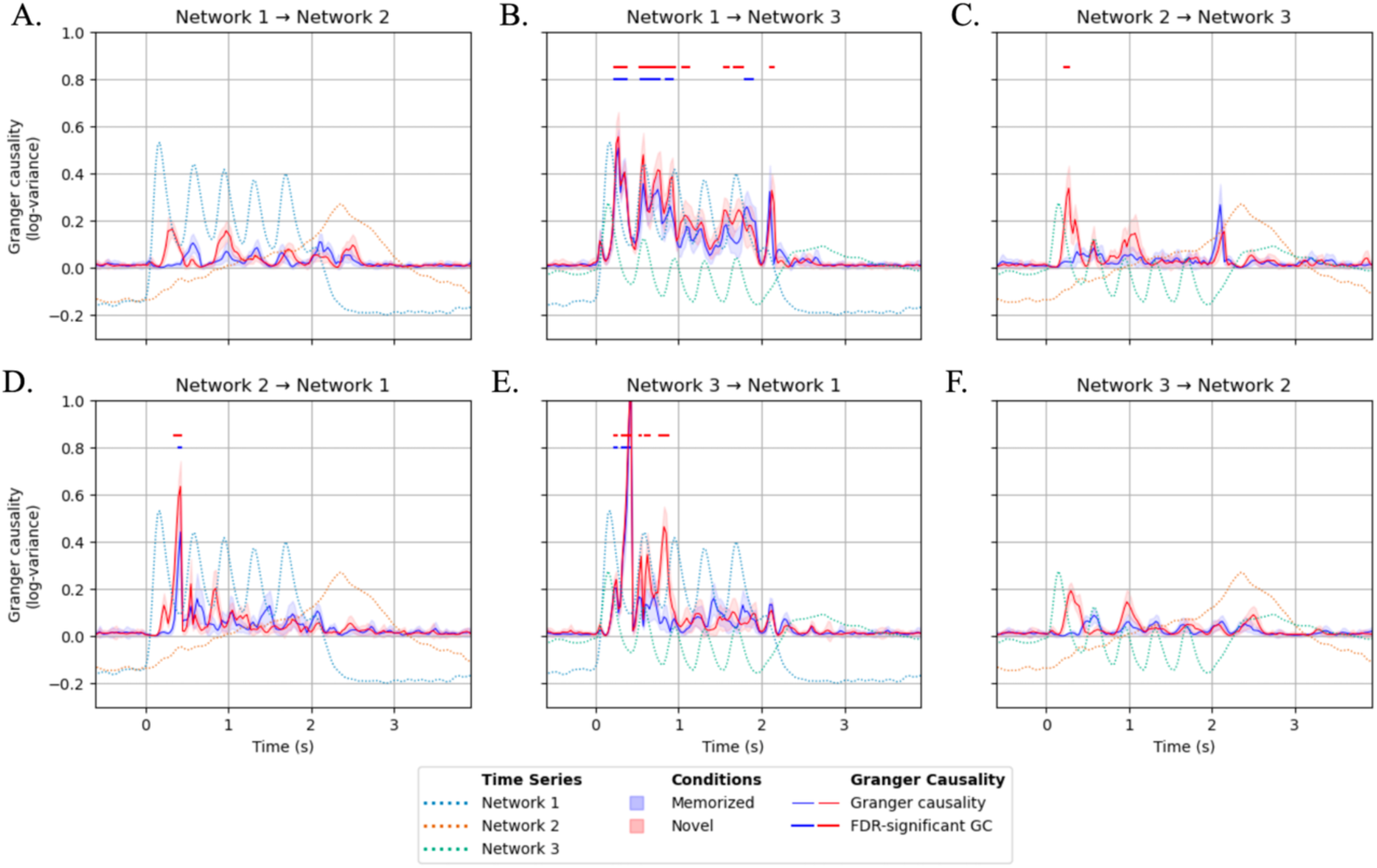
Granger Causality (GC) Mean and standard deviation of GC for each network before the computation of the net GC. Significant directional interactions are indicated by lines above or below the zero net-GC line, according to the direction of information transfer. Statistical significance was determined using surrogate-based empirical p-values, followed by Benjamini–Hochberg false discovery rate (FDR) correction for multiple comparisons across temporal windows (FDR-adjusted p < 0.05). Only significant interactions with an absolute net GC magnitude exceeding 0.2 (|**net GC**| > **0**. **2**) are displayed.

**Figure S6.**
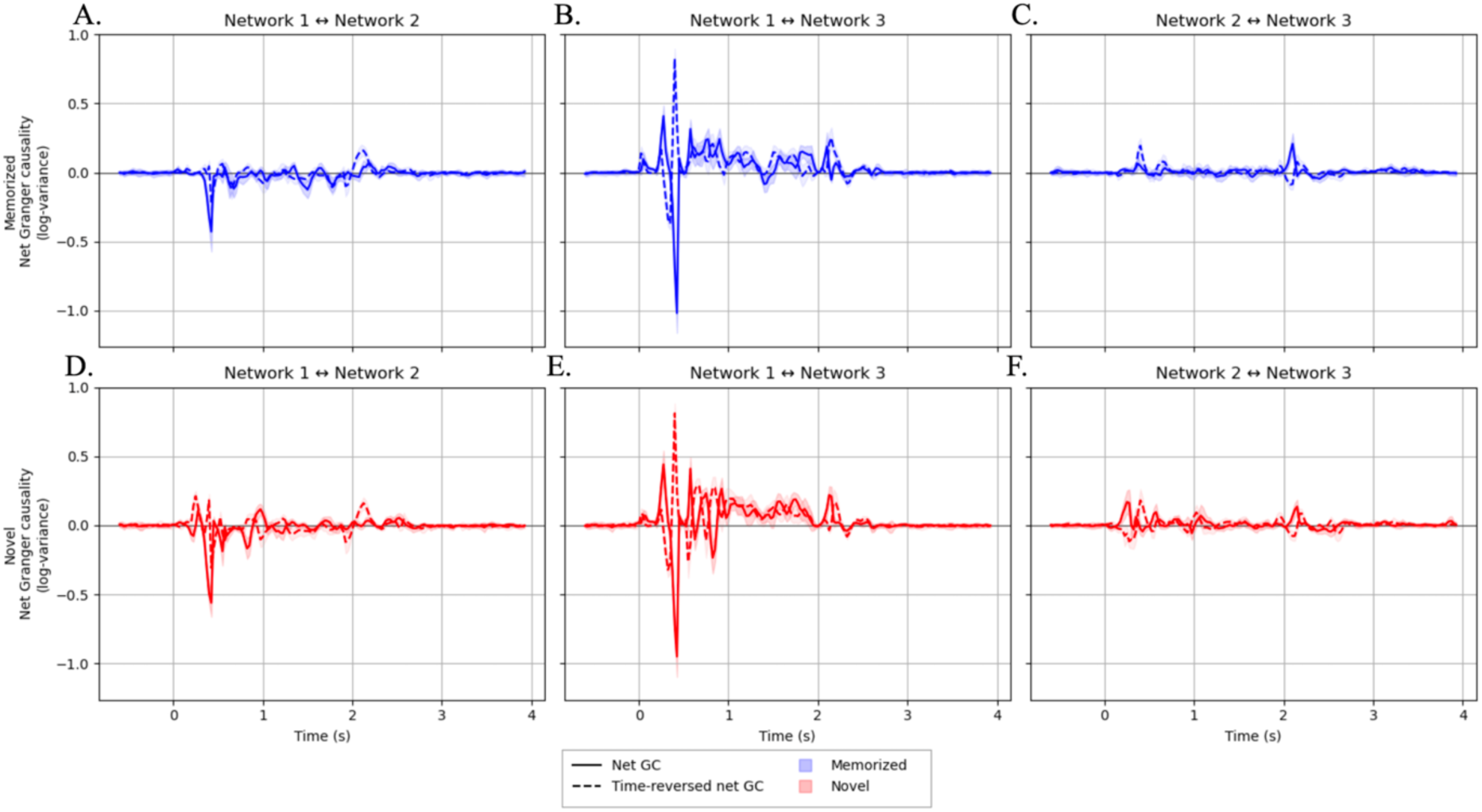
Net Time-reversed Granger Causality Inspection. GC was computed for each network pair using the reversed time sequence and the same method as for non-time-reversed GC. Net time-reversed GC per pair (**a, d-** Network1 and Network2, **b, e-** Network1 and Network3,**c, f-** Network2 and Network3) is compared to Net GC for each condition: **a-c** Memorized, **d-f** Novel. A change in the direction of net GC rules out signal-to-noise directionality confounds.

### Behavioral Correlation with Networks

**Figure S7.**
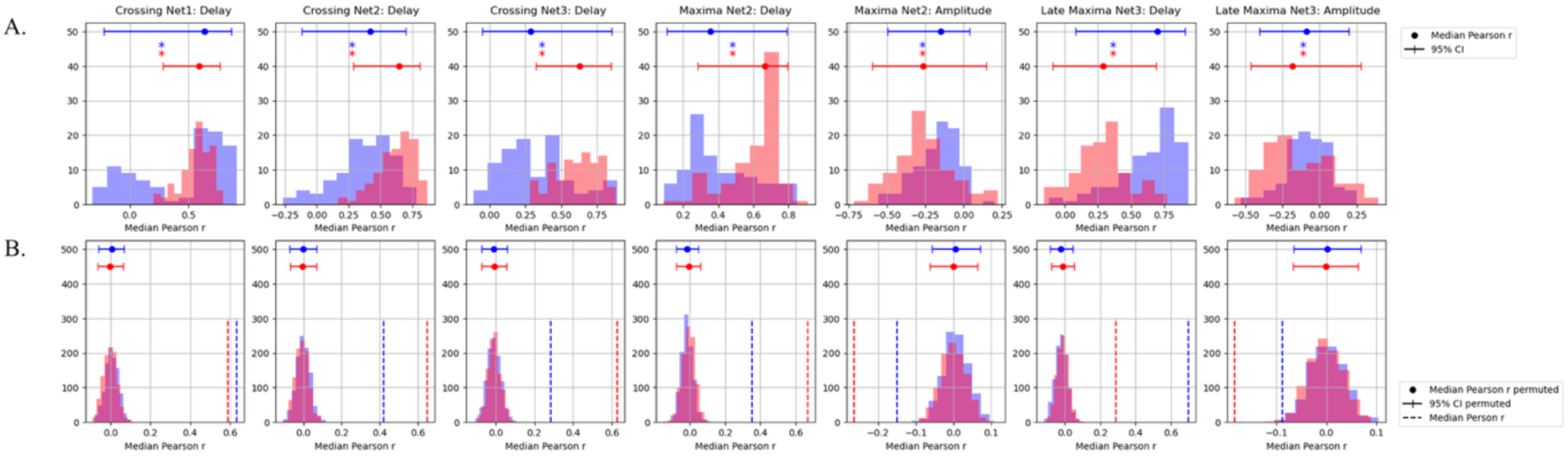
**Motif of interest and Reaction time Pearson correlation.**Pearson correlation between motif features and reaction time mean over runs. Each run corresponds to a randomly subsampled set of trials used to build the “pseudo-subject”. The mean RT across subject trials is used to assign an RT to each resulting “pseudo-subject” trial. **a –** Distribution of the Pearson correlation coefficient between RT and a motif feature across 100 random trial selections and associations. The median and 95% confidence interval (CI) are reported. **b** – The same procedure was performed using shuffled RTs to generate a null distribution (P=1000) for each trial random sampling. Statistical significance was assessed using the median across random sampling to stabilize the correlation. An empirical p-value was computed from the null distribution medians across random sampling.

## Author contributions

Conceptualization: BG, DQM, LB

Writing – original draft: BG

Writing – review & editing: BG, DQM, LB, MR, RTK

Data curation: DQM, AOB, AKS, TE, OKM, JTW, PB, MD, JL, RTK, MR

Investigation: DQM, AOB, AKS, TE, OKM, JTW, PB, MD, JL, RTK, MR

Methodology: BG, DQM, LB

Formal analysis: BG

Software: BG, DQM, MR

Validation: BG, DQM

Visualization: BG

Funding acquisition: DQM, LB

Project administration: DQM, LB

Resources: LB

Supervision: DQM, LB

## Competing interests’ statement

### Code availability

All the code can be found at https://github.com/barbaragrosjean/HG_analysis.git

### Data availability

Data will be openly shared after acceptance in a peer-reviewed journal.

